# Multifractal Analysis of SARS-CoV-2 Coronavirus genomes using the wavelet transform

**DOI:** 10.1101/2020.08.15.252411

**Authors:** Sid-Ali Ouadfeul

## Abstract

In this paper, the 1D Wavelet Transform Modulus Maxima lines (WTMM) method is used to investigate the Long-Range Correlation (LRC) and to estimate the so-called Hurst exponent of 21 isolate RNA sequence downloaded from the NCBI database of patients infected by SARS-CoV-2, Coronavirus, the Knucleotidic, Purine, Pyramidine, Ameno, Keto and GC DNA coding are used. Obtained results show the LRC character in the most sequences; except some sequences where the anti-correlated or the Classical Brownian motion character is observed, demonstrating that the SARS-Cov2 coronavirus undergoes mutation from a country to another or in the same country, they reveals also the complexity and the heterogeneous genome structure organization far from the equilibrium and the self-organization.

## 1 Introduction

Severe Acute Respiratory Syndrome SARS-CoV-2 is a member of the Coronaviridae family causes an illness called COVID-19, which can spread from person to person (Conway, 2020). It has many symptoms such as fever, headache, and tiredness. It causes respiratory difficulties that can cause death, especially for people which health chronic difficulties such as diabetes, arterial hypertension, heart and pneumonic illness. Until now, there is no proven anti-viral or vaccination for the SARS-CoV-2 virus.

Fractal character of nucleic acids distribution in DNA sequences has been widely studied by the scientific community; many papers have been published in literature. Arneodo et al (1996) published a paper deals with the study of the Long-Range Correlation (LRC) character of DNA sequences using the 1D continuous wavelet transform method. Zu-Guo et al (2002) introduced a time series model in a statistical point of view and a visual representation in a geometrical point of view to DNA sequence analysis, they have also used fractal dimension, correlation dimension, the Hurst exponent and the dimension spectrum (multifractal analysis) to discuss problems in this field. Cattani (2010) published a paper deals with the digital complex representation of a DNA sequences and the analysis of existing correlations by wavelets. The symbolic DNA sequence is mapped into a nonlinear time series. By studying this time series the existence of fractal shapes and symmetries will be shown. Eight H1N1 DNA sequences from different locations over the world are analyzed. Audit et al (2001) studied the Long-Range Correlations in Genomic DNA and the signature of the Nucleosomal Structure. Audit et al (2004) published a paper deals with wavelet Analysis of DNA bending profiles reveals structural constraints on the evolution of genomic sequences, Voss (1996) published a paper deals with the evolution of Long-Range fractal correlations and 1*/f* noise in DNA base sequences.

In this paper the 1D Wavelet Transform Modulus Maxima Lines (WTMM) method is used to demonstrate the monofractal behavior of SARS-CoV-2 RNA sequences downloaded from the NCBI database and to estimate the so-called Hurst exponent, the goal is to investigate the LRC character in these sequences. We begin by describing the different DNA coding that will be used.

## 2 Different DNA coding

Many DNA coding of the nucleic acids formed by four different nucleotide which are the Thymine (T), the Guanine (G), the Cytosine (C), the Adenine (A) have been proposed in literature, here we will use the following six coding (Messaoudi et al, 2012):

1. The Knucleotidic DNA coding: T=2, G=-2, A=1, C=-1.
2. Purine coding A=G=1, C=T=-1
3. Pyrimidine C=T=1, A=G=-1.
4. Ameno groupe: A=C=1, G=T=-1.
5. Keto coding G=T=1, A=C=-1.
6. GC coding G=C=1, A=T=-1.

## 3 Wavelet Transform Modulus Maxima lines and LRC in DNA sequences

The 1D Wavelet Transform Modulus Maxima lines (WTMM) method is a wavelet based multifractal analysis formalism introduced by Arneodo et al (1995), the algorithm of the WTMM is composed with five steps which are:

1-Calcultation of the Continuous Wavelet Transform (CWT) which is a function of time or the space and the scale.

2-Calculation of Maxima of the Modulus of the CWT

3-Calculation of the function of Partition *Z*(*q, a*).

4-Estimation of the spectrum of exponents *τ*(*q*).

5-Estimation of the spectrum of singularities *D*(*h*).

For more details about the 1D WTMM method we invite readers to the paper of Arneodo et al (1996) or Ouadfeul and Aliouane (2011).

One of the most important strengths of the WTMM method is the ability to identify the mono or the multifractal behavior of a given fractal process, the linear shape of the spectrum of exponents (*τ*(*q*) = *q* ∗ *H* − 1) is enough to say that a given fractal process is monofractal (*H* is the Hurst exponent). For more details about this ability, we invite readers to the papers of Ouadfeul and Aliouane (2011).

Audit et al (2004) showed that there has been intense discussion about the existence, the nature and the origin of LRC in DNA sequences in last decades. Besides Fourier and autocorrelation analysis, different techniques including mutual information functions, DNA walk representation, Zipf analysis and entropies were used for statistical analysis of DNA sequences. Actually, there were many objective reasons for this somehow controversial situation. Most of the investigations of LRC in DNA sequences were performed using different techniques that all had their own advantages and limitations. They all consisted in measuring power-law behavior of some characteristic quantity, e.g., the fractal dimension of the DNA walk, the scaling exponent of the correlation function or the power-law exponent of the power spectrum. Therefore, in practice, they all faced the same difficulties, namely finite size effects due to the finiteness of the sequence. Authors of this paper demonstrated the necessity of the wavelet transform microscope to study the LRC character of DNA sequences.

Estimated Hurst exponent *H* of the DNA walks using the wavelet transform method is able to say that a given DNA walk is an anti-correlated random walk (*H* <1/2: anti-persistent random walk), or positively correlated (*H* > 1/2: persistent random walk). *H* = 1/2 corresponds to classical Brownian motion (Audit et al, 2004).

## 4 Data analysis and results discussion

In this part, 21 isolate RNA sequence are analyzed using the 1D WTMM method, these sequences are extracted from 21 GenBank downloaded from the National Center for Biotechnology Information (NCBI) database, All these RNA sequences are of peoples infected by SARS-CoV-2 Coronavirus, table 01 shows the code of each GenBank and the origin (Country) of each patient infected by this virus. These RNA profiles are coded using the six coding methods detailed above. Then, the 1D WTMM analysis is applied to these sequences to estimate the so-called Holder exponent for each coded DNA profile. Figure 01 shows the RNA Knucleotidic coding with 512bp as a length, the DNA walk of this sequence is presented in figure 02, the DNA walk at the position n is defined as the sum 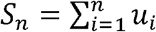 where *u*_*i*_ = {*A*_*i*_, *C*_*i*_, *T*_*i*_, *G*_*i*_}. For more details about DNA Walk, we invite readers to the paper of Peng (1992). To demonstrate the fractal behavior of this DNA walk, we calculate the spectral density |*S*(*f*)|^2^of this sequence and we present the spectral density versus the frequency in the log-log scale (Voss, 1992).It is clear that log(|*S*(*f*)|^2^) has a linear shape (see figure 3), which demonstrating the scale-law behavior of the spectral density versus the frequency 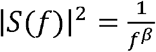, where *β* = 2*H*_*G*_ +1 (*H*_*G*_ is the global regularity, *H*_*G*_ = 0.4) (Arneodo et al, 1995).

**Figure 01.**
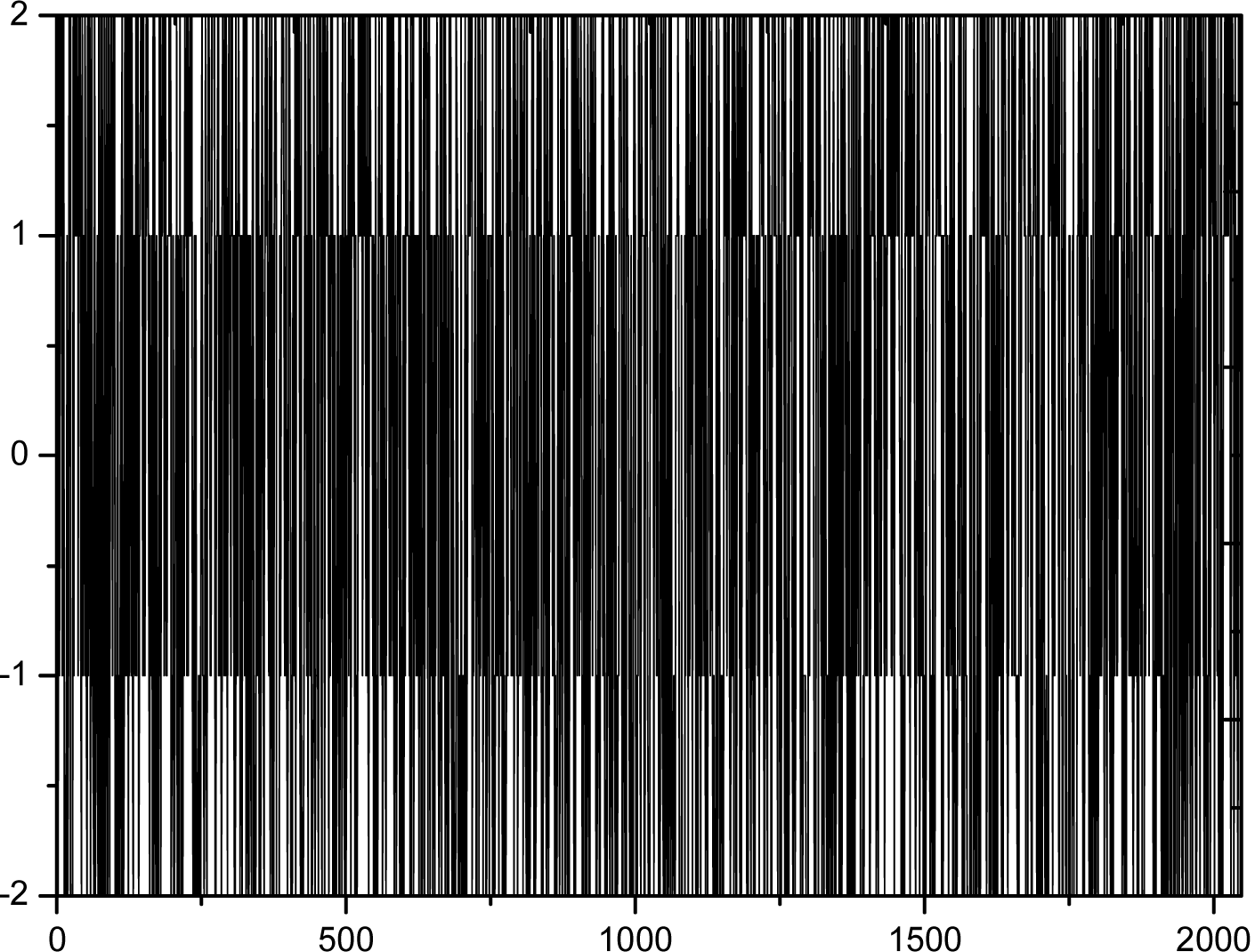
The Knuleoditic RNA Barcoding of the sequence MT066156.1 (Italy) with 512bp as a length.

**Figure 02.**
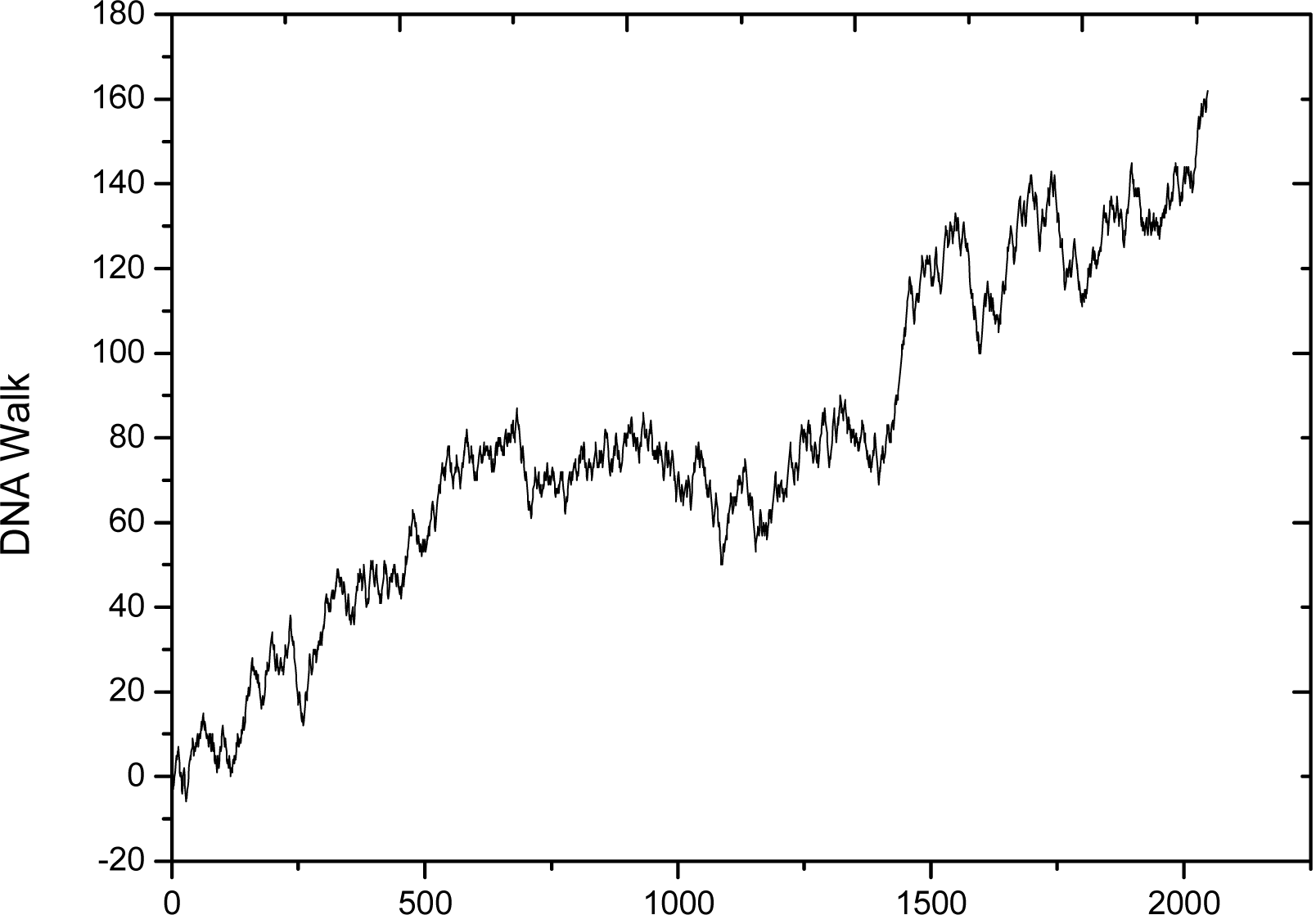
DNA Walk of the sequence MT066156.1 presented in figure 01

**Figure 03.**
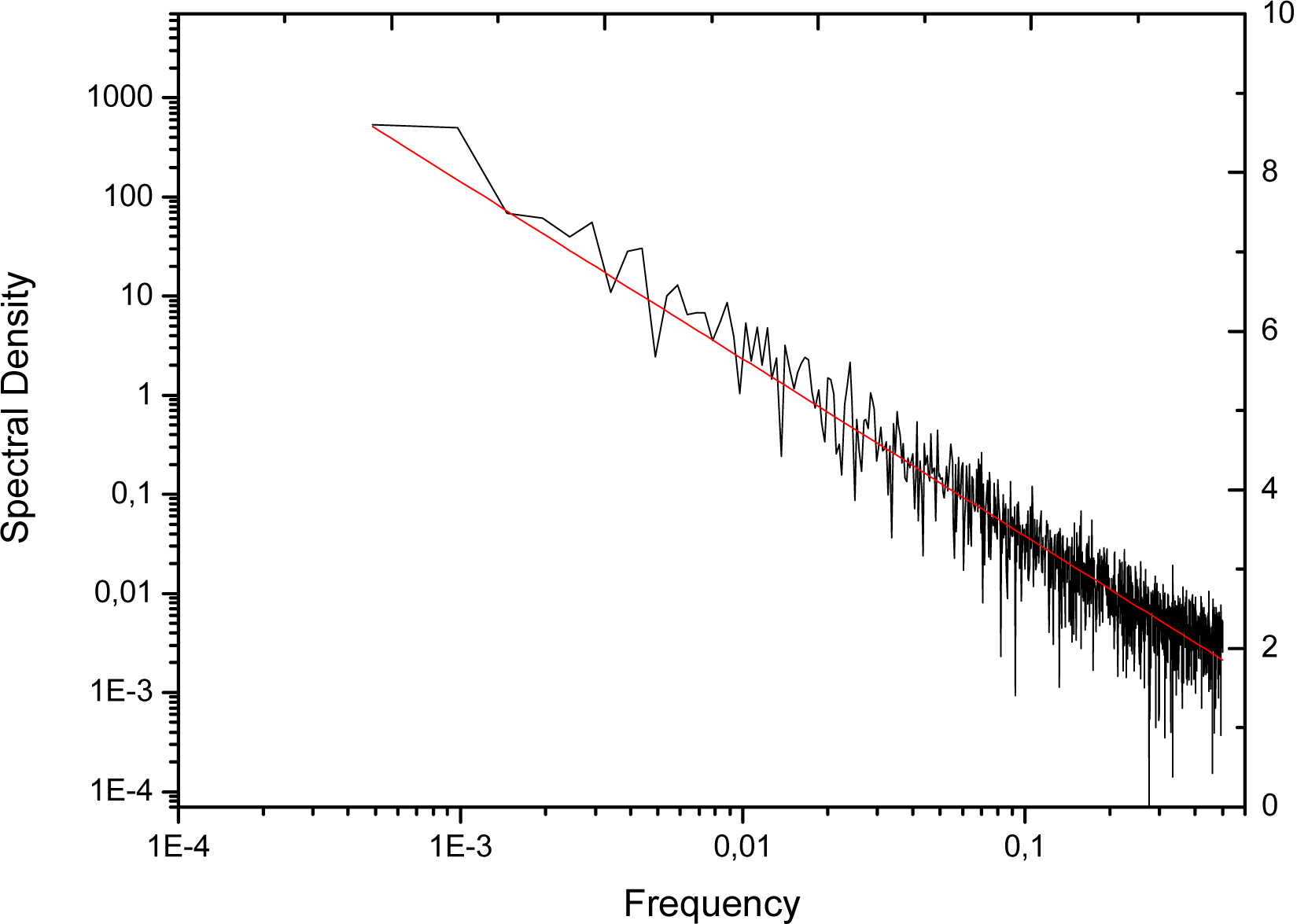
Spectral Density of the DNA walk presented in figure 02

**Table 01.**
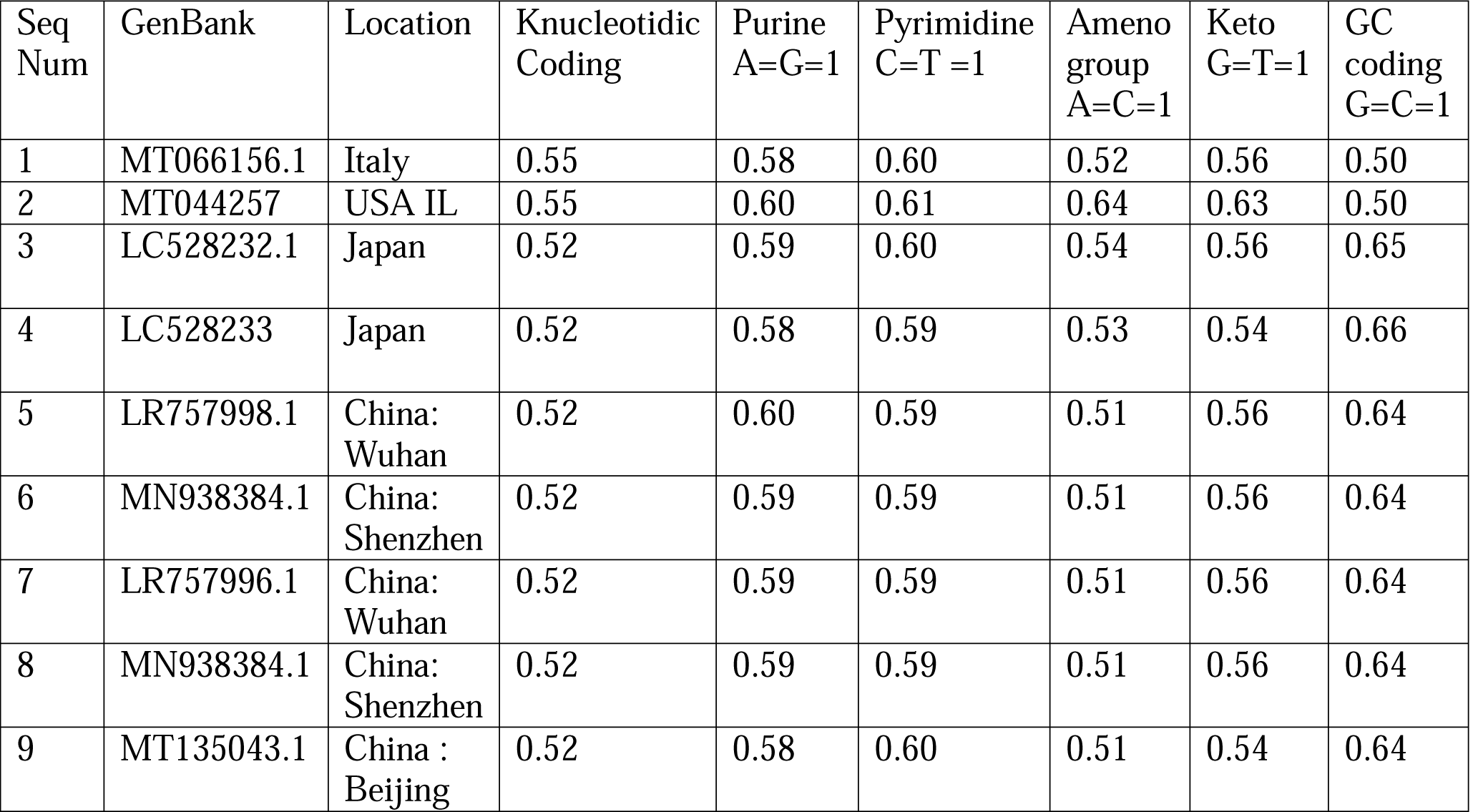
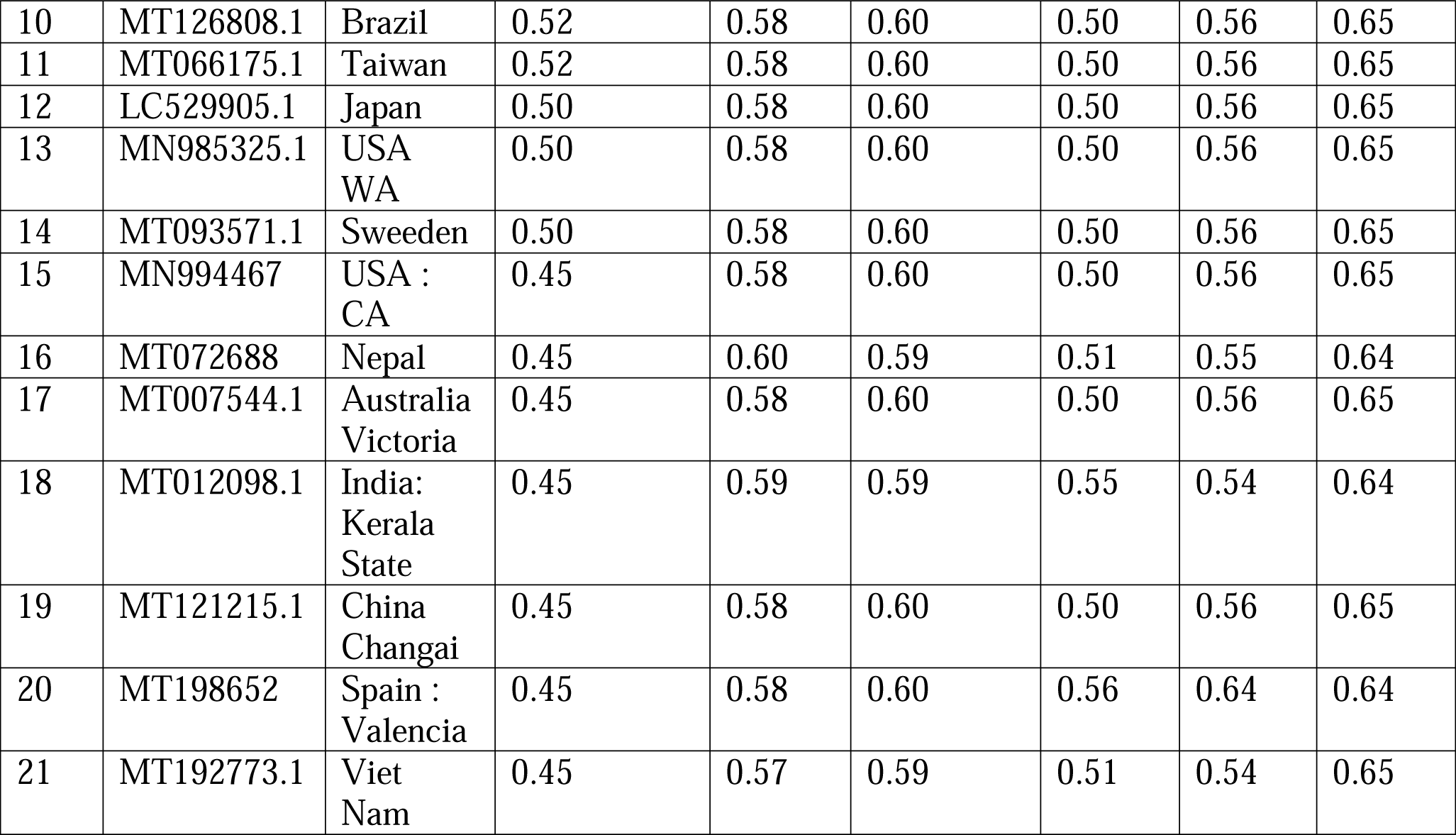
Holder exponents estimated using the 1D WTMM method of 21 Genbank

The first step is the Continuous Wavelet Transform calculation, the analyzing wavelet is the Complex Morlet, for more details about the CWT calculation, parameters of the analyzing wavelet, and the scaling method, we invite readers to the paper of Ouadfeul and Aliouane (2011). Figure 4 shows the modulus of CWT, then maxima of the modulus of the CWT are calculated; graph of chains of maxima is called Skelton, figure 5 exhibits the Skelton of the modulus of the CWT. Then, the function of partition is calculated on the *(q,a)* domain, where *a* is the dilatation and *q* is a scale parameter, for a good WTMM analysis and a high capacity of the WTMM method to identify the mono and multifractal behavior of these RNA coded sequences we take in consideration only the *q* scale parameter interval [-0.5,+0.5] (Ouadfeul and Aliouane, 2011). Figure 6 shows the spectrum of exponents versus *q*, the spectrum of exponents *τ*(*q*) exhibits a linear behavior with equation *τ*(*q*) = *q* ∗ *H* − 1. *H* is the Hurst exponent (slope of the straight line) which is equal to 0.55 for this sequence.

**Figure 4.**
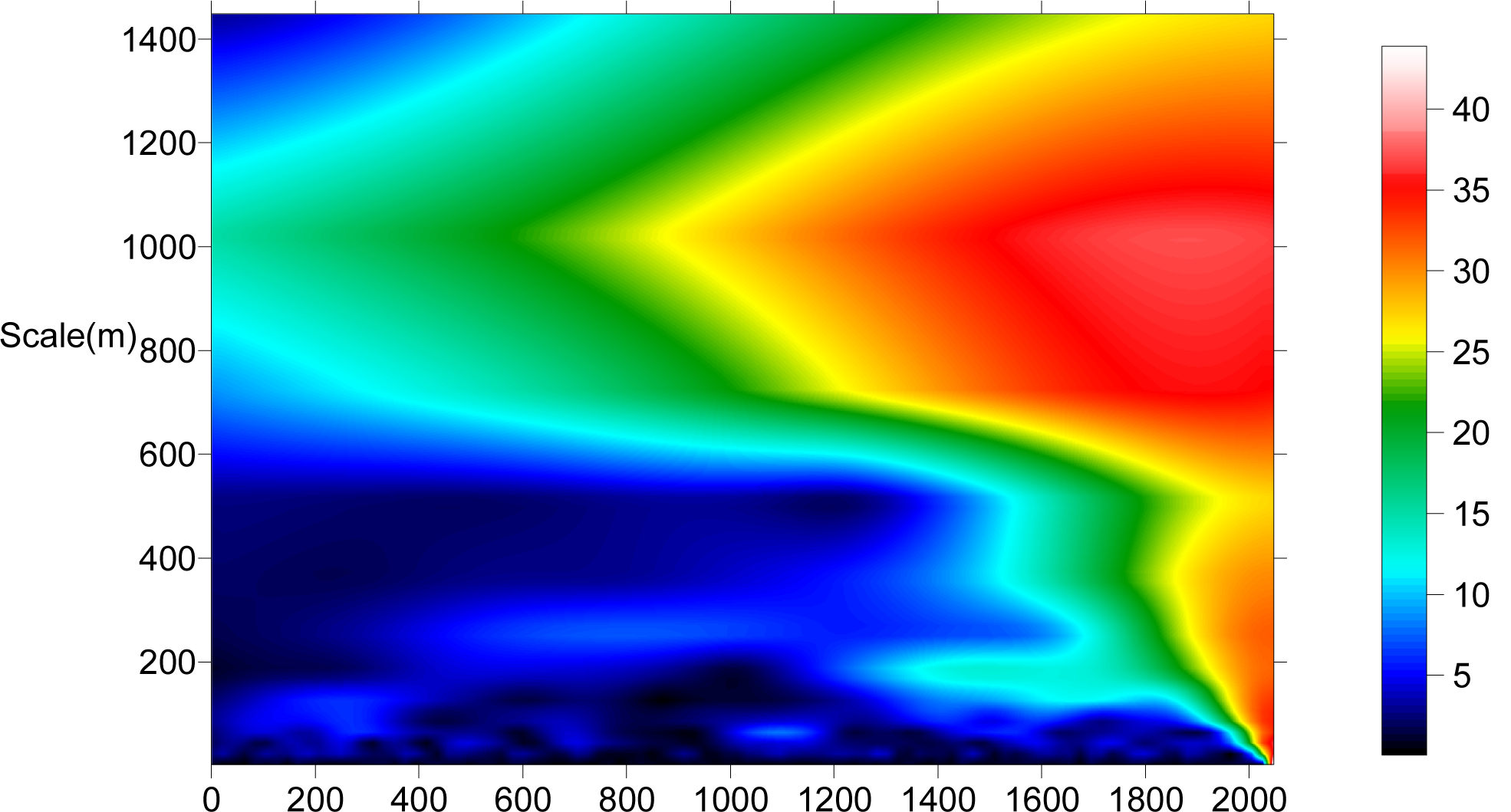
Modulus of the Continuous wavelet transform of the DNA walk shown in figure 2, the analyzing wavelet is Complex Morlet.

**Figure 5.**
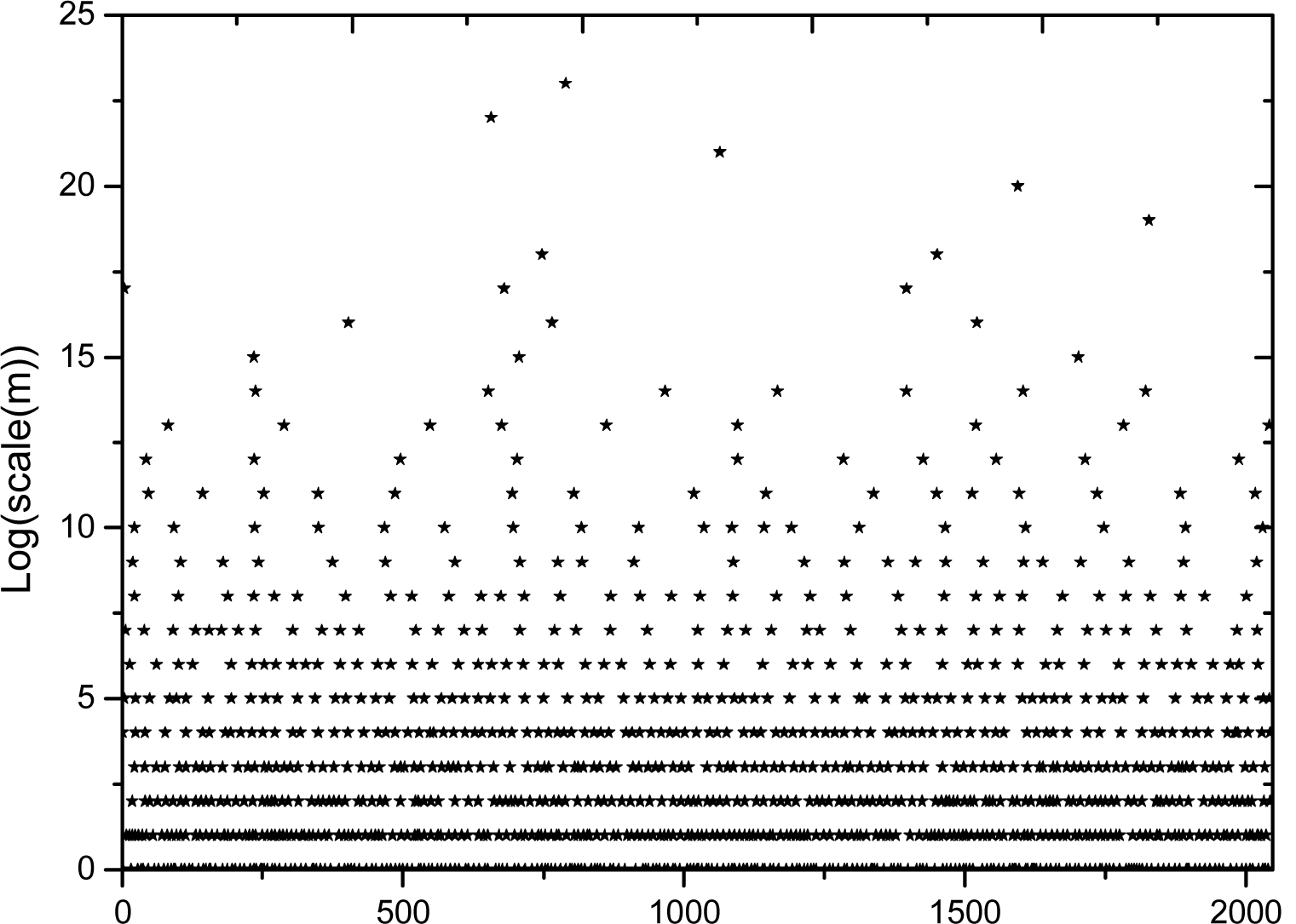
Skelton of the modulus of the CWT

**Figure 6.**
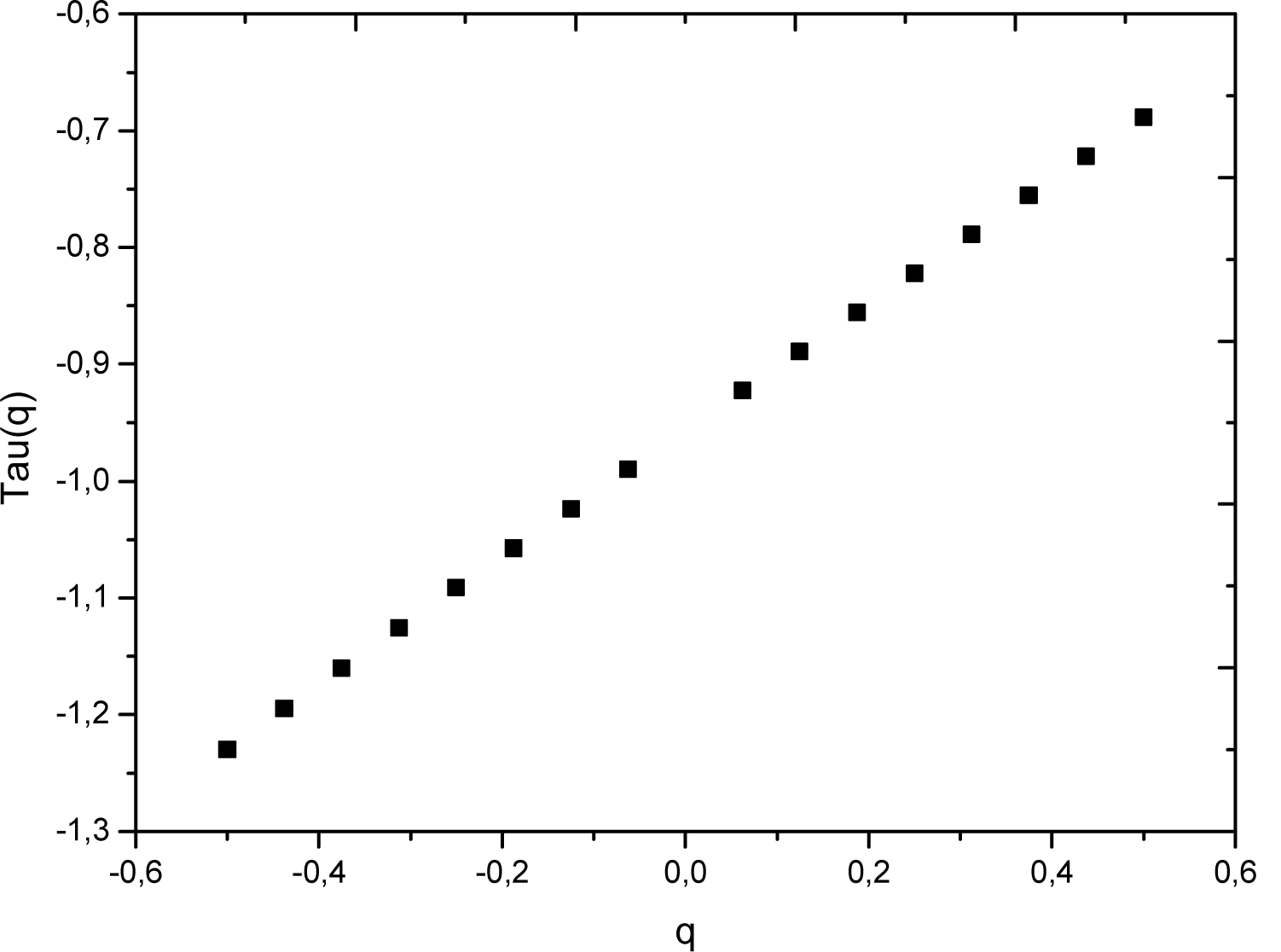
Spectrum of exponents derived from the function of partition

The calculated value of the Hurst exponents demonstrates the mono-fractal behavior of the SARS-CoV-2 RNA sequence (which is in agreement with Audit et al (2004) that have demonstrated the monofractal behavior of human and animal DNA and virus RNA sequences), this value demonstrates also the LRC character of this RNA profile.

Same procedure is applied to the 21 RNA sequences for each RNA coding method, table 01 shows the estimated Hurst exponents, obtained results clearly show the Long-Range Correlation (persistent random walk) for the most RNA sequences except for the Genbanks LC529905.1, MN985325.1, and MT093571.1 (Knucleotidic Coding); MT126808.1, MT066175.1, LC529905.1, MN985325.1, MT093571.1 and MN994467 (Ameno group coding); MT066156.1 and MT044257 (GC coding) where the RNA profiles exhibit a classical Brownian motion behavior (*H* = 0.5). Obtained results demonstrate also that the RNA profiles MN994467, MT072688, MT007544.1, MT012098.1, MT121215.1, MT198652, MT192773.1 exhibit an anti-correlated random walk behavior (anti-persistent random walk) for Knucleotidic Coding. Obtained Hurst exponents by the 1D WTMM method have the same values for the same coding in some countries, for example in Knucleotidic coding the Genbank of the following locations have the same Hurst exponent: (Italy, USA Illinois), (Japan, China: Wuhan, China: Shenzhen, China : Beijing, Brazil, Taiwan); (USA : California, Nepal, Australia Victoria, India: Kerala State, China Changai, Spain : Valencia, Viet Nam) which confirm the possibility of mutation of SARS-CoV-2 virus. Audit et al (2004) debated LRC character in DNA profiles and demonstrated that this character is not a simple reason of text and character replication in DNA sequences but it is due to Bacterial/Viral signature in human/animal cells.

They have demonstrated also that the choice of the DNA coding method is very important to extract the Bacterial/Viral signature.

Cattani (2010) demonstrated that the concentration/replication and hidden symmetries of the four nucleic acids in DNA maps built from DNA profiles and the fractal signature in these maps is a strong indicator of H1N1 viral infection, he has demonstrated also that the wavelet-based fractal analysis of DNA profiles extracted from these maps is also a strong indicator of H1N1 viral infection.

Table 02 shows the average value of the Hurst exponent for each coding method of the 21 RNA GenBank, we can observe that the Purine (sensitive to A and G concentrations) and the Pyrimidine (sensitive to C and T concentrations) have the smallest Hurst exponent variation compared to Ameno (sensitive to A and C concentrations) and GC coding which have the highest variation of the Hurst exponent.

**Table 02.** Average values and variation of the Hurst exponents for the six DNA coding methods.

| DNA coding | Hurst exponent |
| --- | --- |
| Knucleotidic | $0.50 \pm 0.05$ |
| Purine | $0.585 \pm 0.015$ |
| Pyrimidine | $0.60 \pm 0.01$ |
| Ameno | $0.57 \pm 0.07$ |
| Keto | $0.59 \pm 0.05$ |
| GC | $0.58 \pm 0.08$ |

## 5 Conclusions

We have performed a 1D wavelet based multifractal analysis of 21 RNA profiles downloaded from the NCBI database using the continuous wavelet transform, the analyzing wavelet is the complex Morlet, the analyzing parameters of the wavelet transform modulus maxima lines method have been optimized by Ouadfeul and Aliouane (2011) in previous works related to the analysis of geophysical signals. Six coding methods are used; the goal is to investigate the Long-Range Correlation character in RNA SARS-CoV-2 Coronavirus genomes. Obtained results demonstrate the LRC character in the most sequences; except some sequences where the anti-correlated or the Classical Brownian motion characters are observed. Obtained results demonstrate also that the SARS-CoV-2 undergoes mutation, varying from a country to another or varying in the same country like China.

Average value and variations of the Hurst exponent for each coding method of the 21 GenBank reveal the complexity and the heterogeneous genome structure organization far from the equilibrium and the self-organization.

We recommend extending the multifractal analysis to the 2D domain; this can be realized by the use of 2D Wavelet Transform Modulus Maxima lines method which has proven its robustness in many fields and branches of sciences.

## Acknowledgments

Authors would like thank the National Center for Biotechnology Information for providing data.

